# Dopaminergic neurons modulate locomotion in *Caenorhabditis elegans*

**DOI:** 10.1101/056192

**Authors:** Mohamed Abdelhack

## Abstract

Adaptation in the sensory-mechanical loop during locomotion is a powerful mechanism that allows organisms to survive in different conditions and environments. Motile animals need to alter motion patterns in different environments. For example, crocodiles and other animals can walk on solid ground but switch to swimming in water beds. The nematode *Caenorhabditis elegans* also shows adaptability by employing thrashing behaviour in low viscosity media and crawling in high viscosity media. The mechanism that enables this adaptability is an active area of research. It has been attributed previously to neuro-modulation by dopamine and serotonin.

The aim of this study is to physiologically investigate the neuronal mechanisms of modulation of locomotion by dopamine. The results suggest that the mechanosensory properties of the dopaminergic neurons PDE are not limited to touch sensation, but to surrounding environment resistance as well. The significance of such characterization is improving our understanding of dopamine gait switching which gets impaired in Parkinson’s disease.

## Introduction

In nearly all the living organisms, locomotion is an important strategy for survival. It is important for feeding, avoiding predators, and finding optimal environmental conditions for survival. Different species employ different locomotion methods to accommodate different environmental conditions that affect the efficiency of the locomotion process. In order to maximize efficiency of locomotion in different environments and also in different situations (e.g., the presence of a predator), organisms need to have an adaptive strategy that includes a sensory system. Such a system must perceive environmental factors, enabling the organism to modify the locomotion pattern. In robotics, such mechanical sensory feedback loops constitute a growing field of research. There are many factors that can affect locomotion behaviour. For example, mechanical forces imposed by the environment dictate some constraints on locomotion and on how much energy needs to be consumed to overcome the resistance to a moving body. Also, the presence of a predator would evoke an escape response, which is a distinct pattern from normal locomotion in most organisms. Many other factors, such as chemical cues, presence of food…etc, can affect locomotion strategy in various ways. In this study, we focus on environmental mechanical forces that cause an organism to modulate its locomotion pattern in order to optimize energy consumption.

Locomotion is generally a rhythmic process where a sequence of actions is repeated with a certain frequency. This is known as gait. All motile animals exhibit at least one gait which can get modulated. Higher animals exhibit more than one gait where each of them is modulated by the external environment and at certain points gait switching occurs. One example is human walking and running. A human can modulate walking pattern increasing from slow walking to fast walking. To further increase the speed an abrupt change happens where the pattern of locomotion changes from walking to slow running and then running speed can be further modulated. Neuronally, a central pattern generator (CPG) is utilized for generating the locomotion pattern^1, 2^. A central pattern generator is a group of neurons or a neural network that spontaneously generates rhythmic patterns of neuronal activation in the absence of sensory input. All the central pattern generator needs, is a command signal to initiate the pattern and in some species a modulation signal as well. The CPG not only exists in locomotion circuits, but in any rhythmic output pattern circuit. For example, a big portion of our current knowledge of CPGs comes from the crab stomatogastric system^3^. As for locomotion, cats were the first to be studied by preparing a decerebrated cat is placed on a treadmill^4–7^. Rhythmic locomotion was still possible in these cats which led to introducing the concept of the CPG. More detailed studies were conducted on invertebrates like the lobster, crab, crayfish, and leech to discover the neuronal components of CPGs. These systems give the advantage of ease of preparation. The nervous system can be isolated from the physical body and studied individually or it can be partly isolated by keeping parts of the body intact. Electro-physiological recordings from these neurons can be obtained in such settings.

Modulation of the rhythmic pattern of the CPG has been found to be mediated by several mechanisms. In the petropod mollusc swimming circuit, modulation is governed by higher cerebral input^8^ while in crab stomatogastric ganglion, modulation is governed directly by sensory input^9^. Crab pyloric rhythm, on the other hand, is modulated by proprioceptive input from another CPG circuit^10^. In lobster and many other animals, neuromodulators like dopamine and serotonin govern CPG modulation^11^.

*C. elegans* modulates its locomotion pattern. This appears to be a modulation of undulation frequency, undulation wavelength, and velocity of locomotion. There has been a debate as to whether this modulation translates into two distinct locomotion patterns of swimming and crawling^12–14^; or whether it is one locomotion pattern that is continuously modulated^15–18^.

Dopamine is a biogenic amine that has been shown to be associated with the process^14^. In *C. elegans* hermaphrodite, there are 8 dopaminergic neurons distributed into 3 neuron types, namely CEP, ADE, and PDE^19–21^. To date, five dopamine receptors have been identified in *C. elegans*. Two of them are homologous to mammalian D1-type receptors (*dop*-1 and *dop*-4)^22–30^. The other three are homologues to D2-type receptors (*dop*-2, *dop*-3, and *dop*-6)^24, 27, 31–33^. These receptors are widely expressed in *C. elegans* nervous system. The dat-1 gene encodes a dopamine transporter^34–36^, so it is expressed exclusively in all the dopaminergic neurons^35–41^ which is the reason that this promoter was utilized in this study.

Dopamine is responsible for a spectrum of behaviours. It has been shown that it is responsible for slowing of locomotion on the encounter of food^42–47^, in local search behaviour along with serotonin^43^, and in learning and decision making^48–50^. Dopaminergic neurons are known to be mechanosensory^20, 28, 42, 43, 51–53^ that have been shown to respond to harsh touch stimulation^54–56^ with ADE responsible for anterior touch sensation and PDE responsible for its posterior counterparts^57–59^. A study has shown that dopamine is also responsible for gait switching from swimming to crawling^14^. They have also shown that serotonin is important and sufficient to switch from crawling to swimming. The dopaminergic neurons shown to be responsible for this switching were ADE and PDE but not CEP.

In humans, gait dysfunction is a main symptom of the Parkinson’s disease. It causes failure to modulate locomotion pattern by gait switching^60^. Other symptoms include trembling, stiffness, slowness of movement, and walking difficulty. It is caused by death of dopaminergic neurons in the substantia nigra^61^. The cause of death of these cells is so far unclear^62^.

In this study, I measure the calcium activity of dopaminergic neurons PDE as the worms switch between crawling and swimming due to crossing a viscosity separation to demonstrate that the worms respond to environment sensation by change in basal activity level and show the bidirectional nature of the dopamine effect. While the previous study^14^ has shown that dopamine is responsible for switching from crawling to swimming as dopaminergic neurons are active at switching. Here, I show that dopaminergic neurons also respond to the environment as the worm is switching from crawling to swimming by a decrease in their basal activation which further supports the idea that dopaminergic neurons sense the pressure induced by the environment. I limit the investigation to the PDE neurons only because of the reliability of the extracted signals from these neurons compared with the ADE neurons that are located in the head region which moves faster than the body region where PDE neurons are located and thus causes ADE’s fluorescence signals to be much more ambiguous. In addition, ADE neurons are located close to the CEP neurons which makes signal contamination easier. The investigation is also limited to the transition from high to low viscosity as the opposite has already been investigated^14^, and also because of the impossibility of the worm crossing spontaneously from low to high viscosity in the presented experimental setting due to the insufficiency of thrust.

## Results and Discussion

### Dopaminergic neurons sense the physical environment

As the worms cross the viscosity separation (Figure 1), PDE neurons have shown deactivation responses (Figure 2). The deactivation response is shown not to be a result of either photo-bleaching or loss of focus due to z-axis movement as both controls show very small decrease in activation compared to the neuronal signal of the worm crossing the viscosity separation. Frequency of head undulation of *C. elegans* increases accordingly, however even if it does not increase to reach the swimming typical frequency (Figure 2b), still the corresponding PDE response goes down. This suggests that PDE neurons respond by deactivation to the environmental pressure and not to the body bends. However, they still showed response to body bends which was smoothed out.

**Figure 1.**
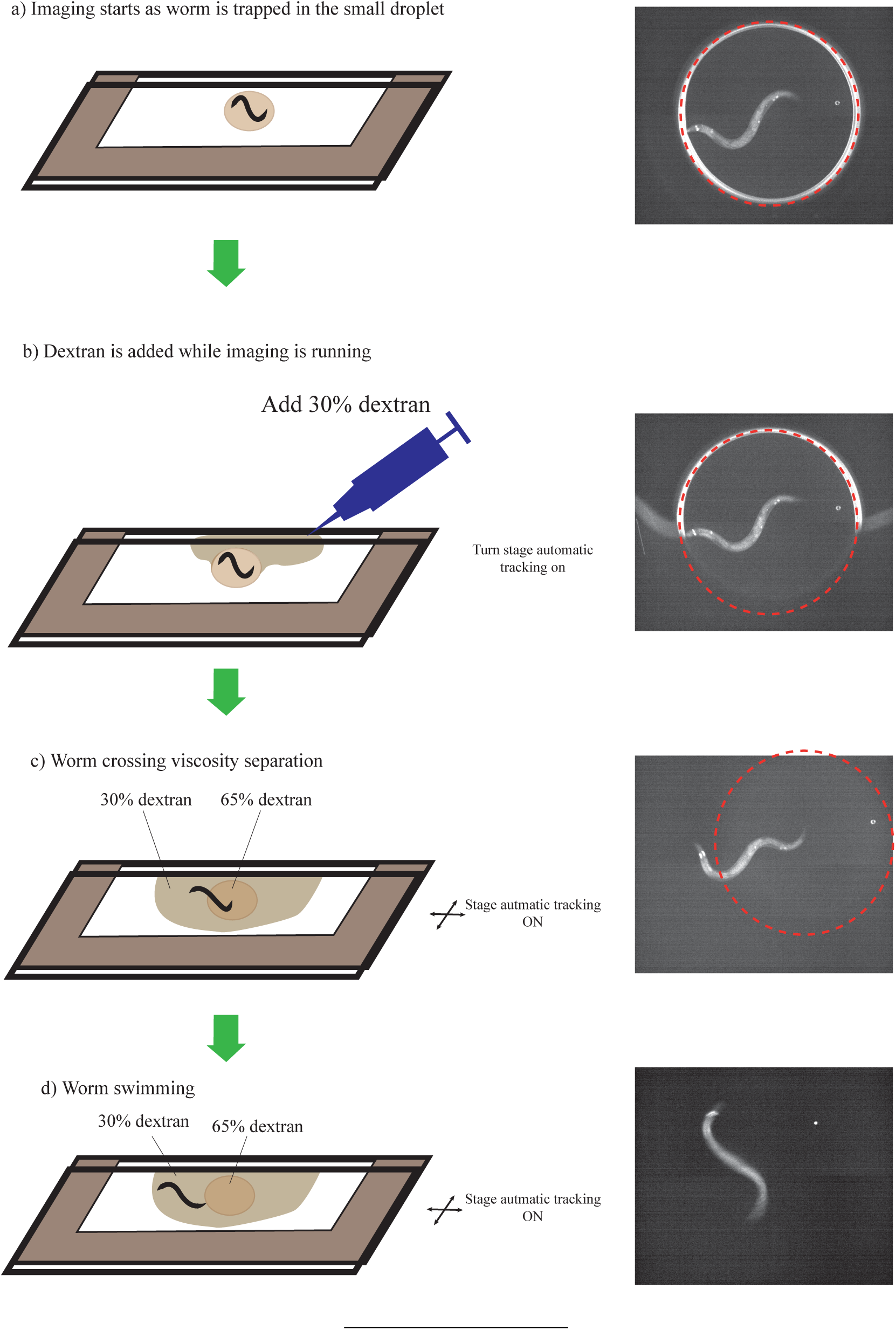
A description of the steps of the viscosity separation experimental protocol where the worm is initially confined in a small droplet of 65% dextran (a) and then dextran 30% is added with micropipette (zero time point) (b). The worm then starts to move out of the small droplet (c) until the whole body gets to the lower viscosity and swimming is maintained (d).

The mean calcium response to crossing the separation is shown in figure 3 where zero time point refers to the time when 30% dextran is visible to have surrounded half of the 65% dextran droplet (Figure 1b). I show also response of up to 70 seconds after the zero time point since the worm’s body, and hence the PDE neurons, takes 10-60 seconds to fully move from the 65% dextran and be fully swimming in 30% dextran which is apparent in the gradual increase in undulation frequency (Figure 3b). The transition time is variable among worms, so the mean calcium signal shows gradual change until it stabilizes at about 60 seconds to about 75% of its value in high viscosity. In some cases we observe a sudden drop while in other cases the drop is gradual (Figure 2). This depends on the speed the worm crosses the separation which is variable. When the worm takes more time to pass the gradient, the sensillar endings of PDE neurons move through the viscosity gradient more slowly and hence the calcium response goes down more gradually. The separation described is also meant to be a very steep gradient of viscosity but as time passes and due to diffusion and stirring effect caused by the worm movement, the gradient can get smoother and hence the transient in the physical forces sensed gets smoother. This can explain why the fluorescence signal drop is more gradual than expected in some cases. For this reason, in figure 3a, the mean signal of the PDE neurons is close in level to the controls until the last 10 seconds when all the worms have moved completely to the low viscosity medium. This could also be attributed to the slow response of these neurons. The ADE neurons, which are similar in genetic identity to the PDE neurons, have been shown to exhibit slow response to the harsh touch stimulation^58^. The signals from 60 to 70 seconds were compared to signals from 10 seconds before the zero time point and are significantly different (Figure 4) (p<0.001, Unpaired two-sample t-test) where frequency of undulation, as expected, is also significantly different (p<0.001, Unpaired two-sample t-test).

**Figure 2.**
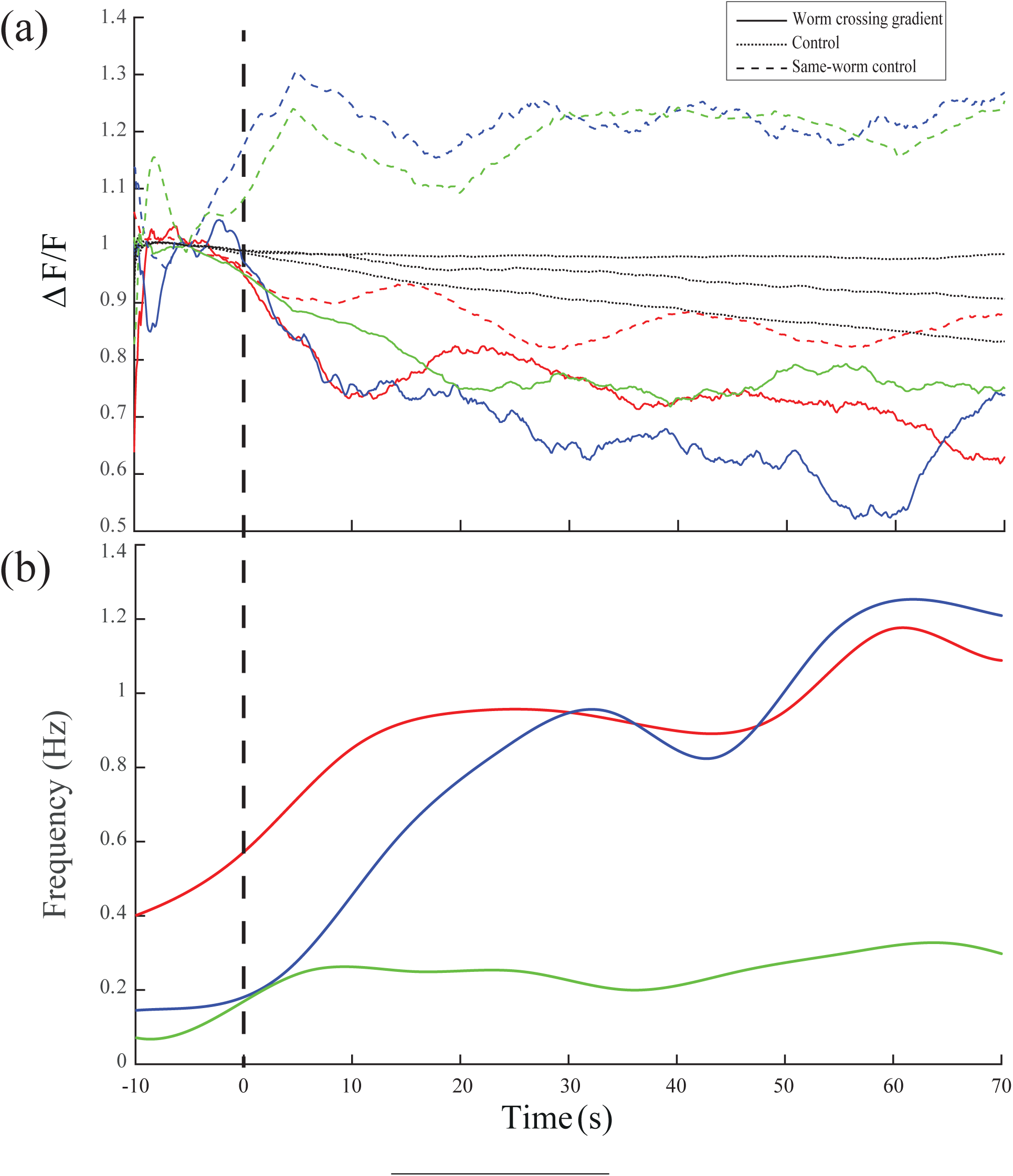
PDE neurons and frequency of undulation response to crossing viscosity separation for three sample worms. Solid lines represent neuronal fluorescence signals of PDE neurons of the worms that cross the separation while dashed lines represent the control signal with mid-body auto-fluorescence signal for the same three worms. Matching colors denote same worm sample. Dotted lines represent three control worms which remain at high viscosity. Since control worm samples are different so they are not color coded. (b) Undulation frequency responses of the same three worms in (a) as the worms switch from crawling to swimming while crossing the viscosity separation. The frequency was measured each 10 seconds and then linear interpolation was done to get the smooth curve. Vertical dashed line denote the zero time point.

**Figure 3.**
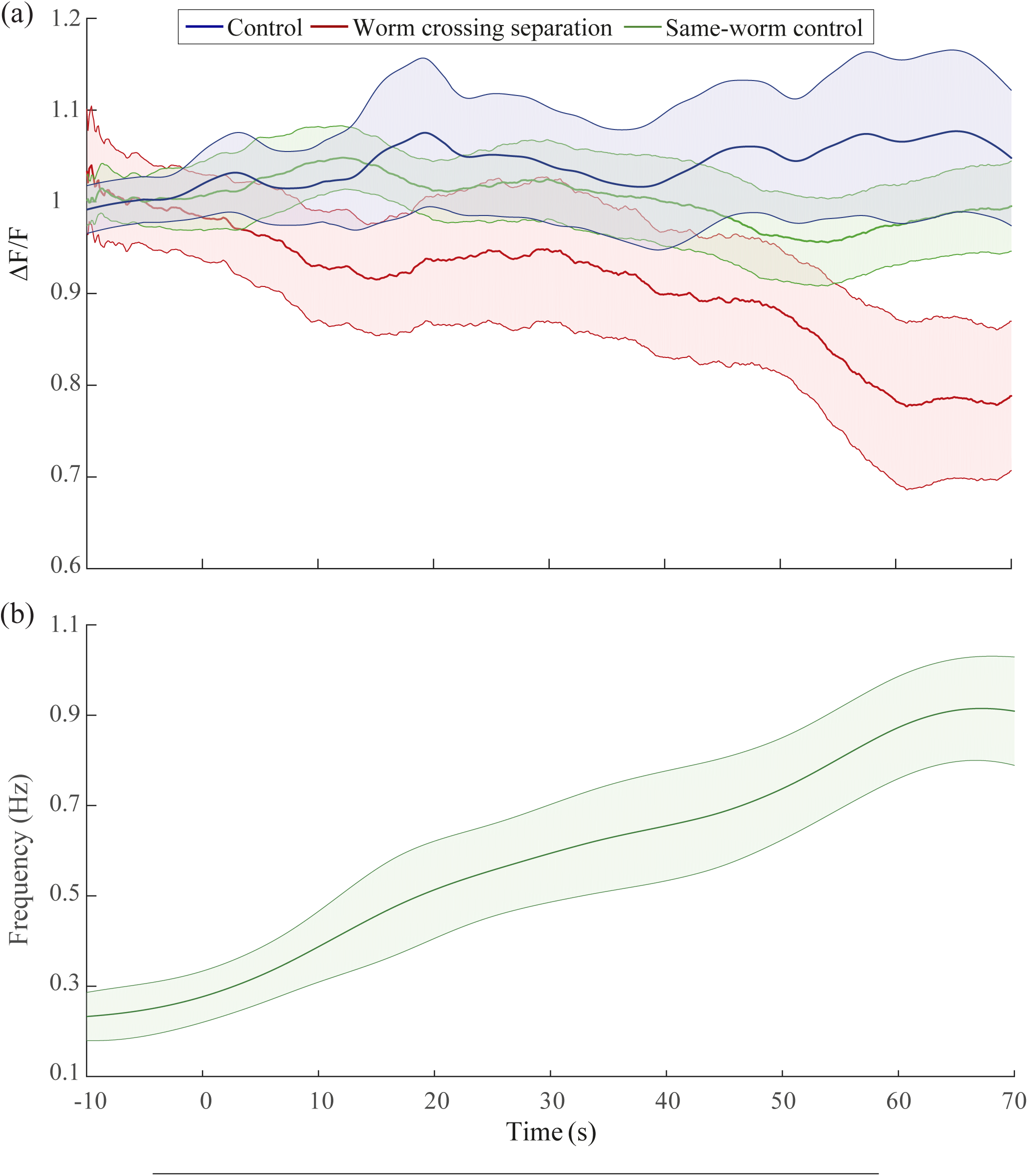
Mean fluorescence signal of PDE neurons and corresponding behavioural response to the worm crossing viscosity separation: (a) Average PDE response to crossing the separation where the zero point is the point of addition of the droplet of liquid (n=10). The control represents worms that remain at high viscosity (n=10). Same-worm control is the measurement of auto-fluorescence signal from the mid-body in order to ensure that the measured decrease in activation is not due to loss of focus. (b) Average frequency response to crossing the separation of the worms in (a). Error bars correspond to ±2 SE.

The transition period could have unexpected signal changes due to many reasons including z-axis movement and stirring effects. Thus, it was important to compare the signal level between worms undulating in homogeneous media. Hence, the mean difference between the PDE signals from 60 to 70 seconds and signals from 10 seconds before the zero time point was compared for the PDE neurons crossing the separation and across the two controls (Figure 4c) The PDE neurons’ signals are shown to decrease by 21.4% on average after crossing the viscosity gradient compared to its signal in high viscosity medium. This decrease is significant compared to a decrease of 1.7% in the same-worm auto-fluorescence signal (p=0.0211, Mann-Whitney U test) and to an increase of 6.3% in the worm in high viscosity neuronal signal (p=0.0211, Mann-Whitney U test). On the other hand, the signal change compared between the controls did not show any statistical significant difference from each other (p=0.7337, Mann-Whitney U test) indicating either minimal z-axis movement or no effect on signals collected using the widefield fluorescence microscopy due to z-axis movements. This indicates the success of the used procedure in measuring the physiological change in the PDE neurons due to crossing the viscosity separation.

**Figure 4.**
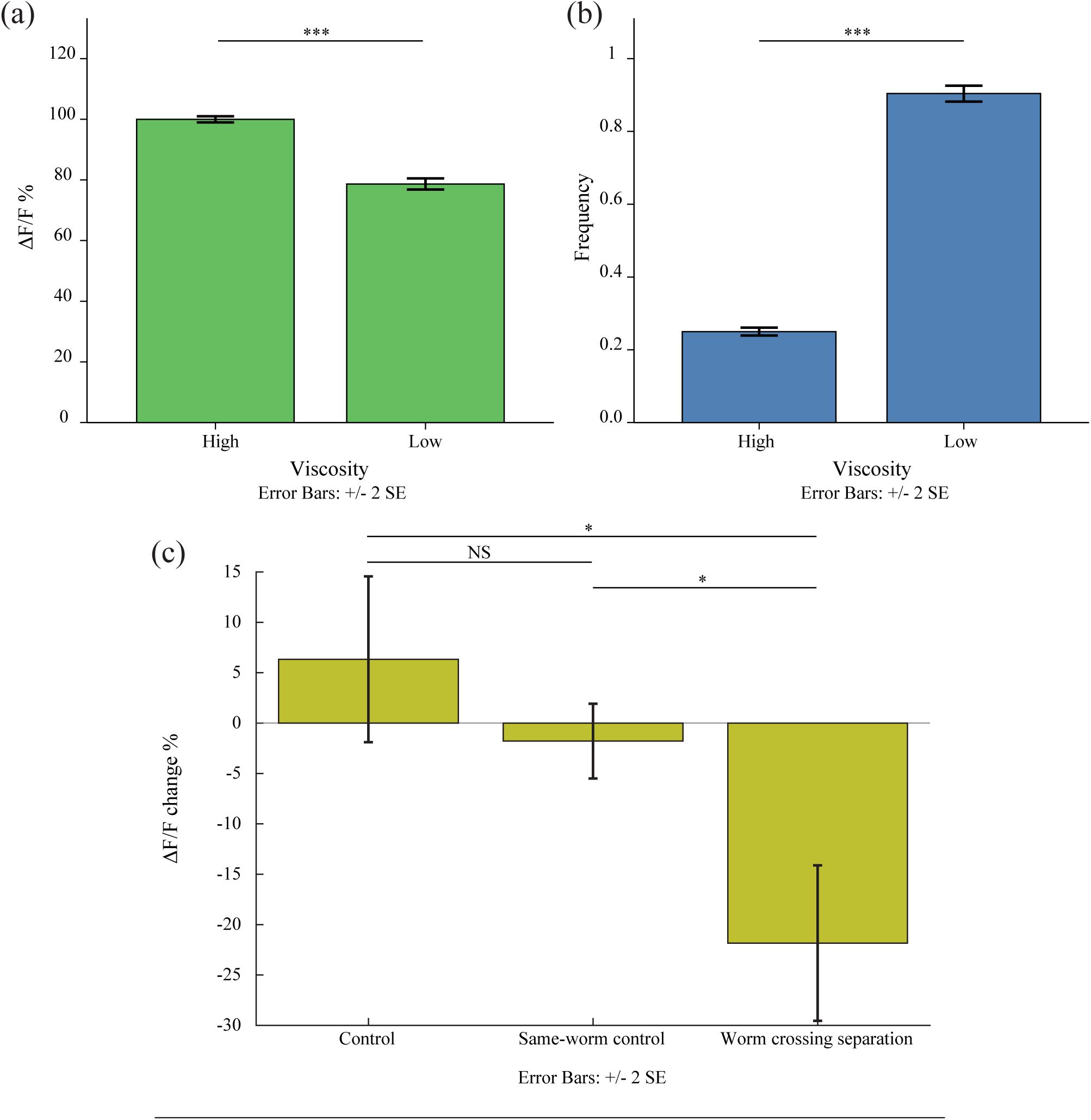
Comparison of mean of data from 10 seconds before the zero time point and from 60-70 seconds after zero time point where the worm in each case is in a completely homogeneous viscosity. (a) Mean activation of PDE neurons in high and low viscosities showing significant difference. (b) Mean frequency of undulation that also shows two different frequencies that are characteristic of both crawling in the high viscosity case and of swimming in the low viscosity case. (c) The mean difference of fluorescence signal of the PDE neurons in the worm crossing separation in comparison to the mid-body auto-fluorescence signal and to the PDE neurons signal from the control worms that remain in the high viscosity medium. It shows that the difference in the observed decrease in activation is a result of the physiological change associated with sensing the environment and not to photo bleaching or loss of focus.

### Calcium signal of PDE and frequency of undulation correlate negatively

In order to find a relation between frequency of undulation and calcium signal, the unsmoothed calcium signals were plotted against frequency of undulation. First, paired-sample t-test was performed on the data and the null hypothesis was rejected (p<0.001). Linear fitting was done by fitting a first degree polynomial *f*(*x*) = *px* + *c* where *p* is the correlation coefficient. The fit showed a correlation coefficient (*p* = -0.2419) Goodness of fit was estimated to be r^2^=0.1232 which is low due to fitting of all data together (Figure 5). This causes the observed non-uniform distribution of the data since the fluorescence levels are expected to be different in different animals due to variability in expression of the vector in transgenic animals. However, they all show the same trend of change. The same polynomial fit was attempted on each time series which shows better results since signals are more self-consistent. Results of fits for each sample are shown in table 1. Another reason for the non-uniformity of distribution is the movement artefacts caused by the neuron sensing body contractions since the dataset used in the fit is unsmoothed. Some vertical clusters and diagonal lines are observed also in the dataset in figure 5. These are caused by the inpainting process that is done to fill in the gaps in fluorescence levels data caused by the lack of reliability of measurement mentioned in the methods section. These interpolations do no include noise from movement artefacts which causes them to form the diagonal lines. Spline interpolation is also done to fill in the gaps in frequency measurements which is measured every 2 seconds. Spline interpolation by nature causes slope of the interpolated curve to be low around real data points which causes the measurements to cluster around the real values and form what can look like vertical lines.

**Figure 5.**
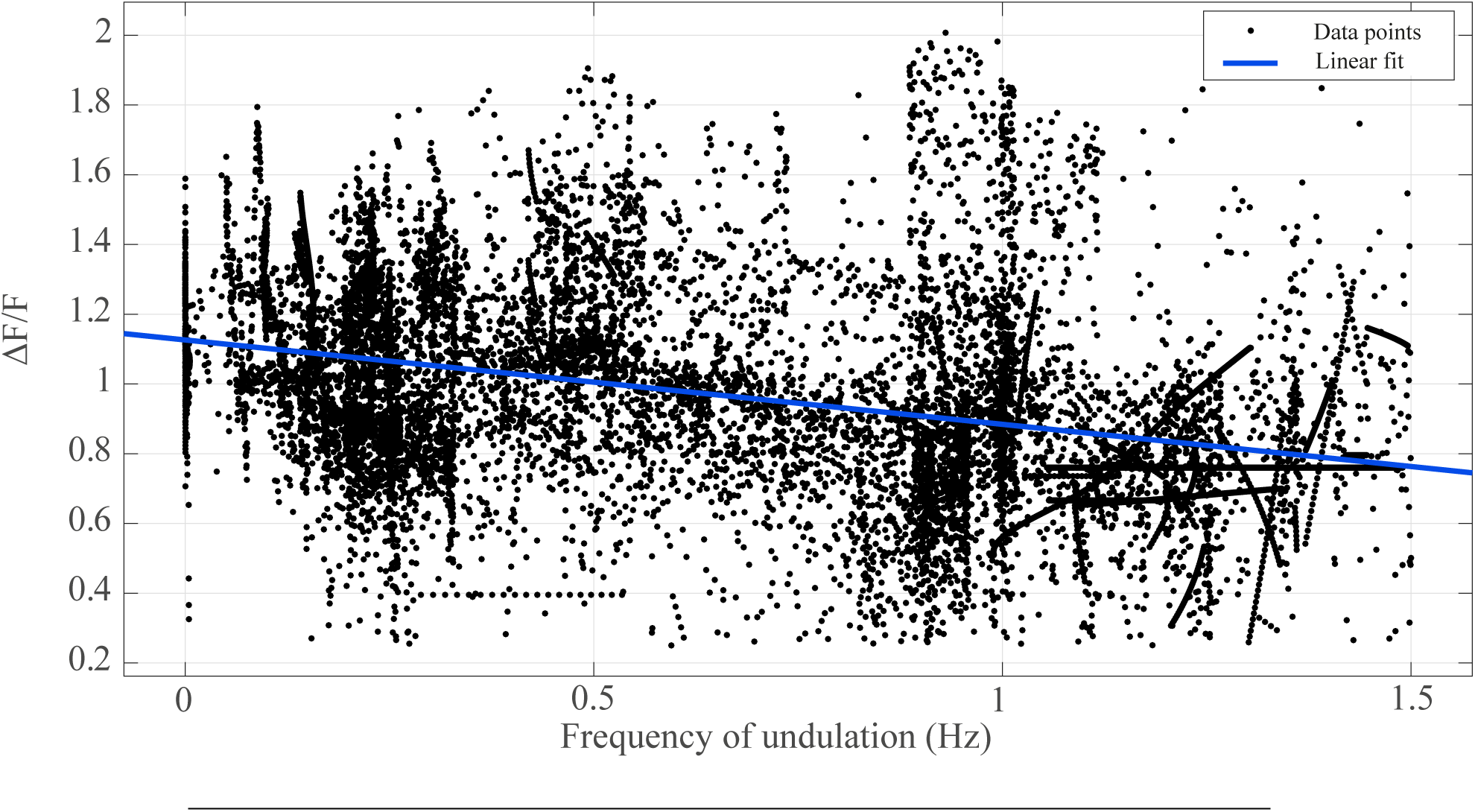
Mathematically fitting the data from fluorescence intensity difference in PDE neurons (y-axis) with the frequency of undulation (x-axis). This figure attempts to fit all the normalized yet unsmoothed raw data of different worms together which leads to the low fitting performance. The detailed fitting results of each sample are shown in table 1.

**Table 1.** Linear fit results of fluorescence intensity difference vs. undulation frequency for each individual worm. In the weighted average, each value of *p* is weighted by the goodness of fit (r^2^) of the same fit, while in the case of average, equal weights are applied. Overall data is the result of fitting all data together shown in figure 5. Only one worm shows a positive *p* coefficient value, but the goodness of fit value is very low rendering data from this sample an outlier.

| Worm | $p$ | $r^2$ |
| --- | --- | --- |
| 1 | -0.01745 | 0.174 |
| 2 | -0.02725 | 0.4107 |
| 3 | -0.01693 | 0.09379 |
| 4 | -0.05399 | 0.4188 |
| 5 | 0.0125 | 0.008285 |
| 6 | -0.0262 | 0.2099 |
| 7 | -0.04159 | 0.1135 |
| 8 | -0.02897 | 0.3299 |
| 9 | -0.01399 | 0.0838 |
| 10 | -0.02735 | 0.1083 |
| Average | -0.0241 | 0.1957 |
| Weighted Average | -0.0318 | N/A |
| Overall | -0.2419 | 0.1232 |

### Hypothesized circuit for forward locomotion

Modulation of undulation frequency observed goes in agreement with the previous reports^12, 14, 16, 17^. The role of PDE neurons in locomotion modulation was further confirmed by neuron activity measurement. A decrease in the neurons activation follows the decrease of external drag imposed by medium viscosity. This can imply a decrease in dopamine release. Another experiment to show an increase in activation level upon switching from swimming to crawling was attempted. The experiment was not successful because in the mentioned experimental setting for viscosity separation, the worm failed to leave the low viscosity medium even after adding an attractant in the high viscosity medium. Moreover, previous reports have shown that activation of dopaminergic neurons causes decrease in speed of locomotion and switching from swimming to crawling^14^. Here I show that they are also involved in switching from crawling to swimming by a decrease in their basal activation. It has been shown also that activation of serotonergic neurons induce increase in head bending frequency that is characteristic of swimming, so serotonin was implied to cause switch from crawling to swimming. Here, it is shown that a decrease in dopamine level is also associated which further supports the results from Topper 2013 where he inhibited *dop*-1 and *dop*-4 expressing neurons which increase time to crawl onset^63^. Thus, it can be suggested that the balance between dopamine and serotonin levels is responsible for mediating crawling-swimming switching. This can be further tested by simultaneous imaging of activation level of both dopaminergic and serotonergic neurons. In a previous study, Topper has found that RID, RIS, and PQR are the neurons responsible for dopamine-mediated gait switching from swimming to crawling^63^. RID and RIS also express serotonin receptors *ser*-2 and *ser*-4 respectively so it can be suggested that they are the ones that compute the dopamine-serotonin balance and modulate locomotion accordingly. RIS also synapses to the head motor neurons RMD and has been shown to inhibit head movement during sleep-like state^64^ where dopamine has been shown to have a role in^65^. RID has gap junctions with the forward locomotion command interneurons AVB which in-turn activate the motor neurons in the body region. Calcium imaging of these two neurons would be a logical next step.

The study also shows that mechanosensation has an effect on normal locomotion behaviour agreeing with a previous report^54^. This is the first report to attribute such role to PDE neurons through a direct measurement of its activity. I show that viscosity differences can be measured by mechansensory neurons and it yields significant change in basal activation level of the neurons. The lower basal activation level during swimming can suggest an explanation to the touch insensitivity in swimming worms compared to crawling worms. As for PDE, the basal activation level is lower during swimming. To reach a significant level enough to induce touch escape response, the mechanical stimulus has to be stronger to compensate for the difference between swimming and crawling activation levels and reach the threshold for escape response.

## Methods

### Behavioural assay

Young adult *dat-1p*::*GCaMP3* worms^58^ (Gift from Prof. Dr. David Biron, The University of Chicago) are placed between two glass slides with a paper separator of thickness 0.1-0.2 mm to minimize movement along the z-axis (n=10). The paper is coated with grease to secure the slides together (Figure 1). The choice of these dimensions will cause the gap between the two plates and the body of the worms to be less than the body diameter. Thus, the wall effects on th normal drag forces will always push the worms to the mid-point between the two plates keeping it always in focus of the objective lens^66^ (except while crossing the gradient where the wall effect would be difficult to calculate). Worms are moved to a foodless plate before assay while the glass cassette is prepared. A tiny droplet of 65% dextran is placed on a glass slide so that the whole droplet is visible under microscope’s field of view. The worm is picked and added to this droplet by a platinum wire worm pick. The second glass slide is then placed on top with the separator in the middle. The top plate is sheared by some distance from the lower plate to ease addition of the low viscosity medium later. Imaging starts while the worm is confined in the tiny droplet (Figure 1a). After the imaging starts, 30% dextran is added with a micropipette to fill the area around the 65% dextran droplet and then tracking starts (Figure 1b,c). The worm immediately starts moving towards the 30% dextran medium to avoid its confinement and because of the lower resistance. Imaging continues until the whole worm’s body passes through the viscosity separation to the lower viscosity region (Figure 1d). The time taken for the whole worm to fully transfer from the high viscosity medium to the lower viscosity one is variable. However, in all samples presented here, before the last ten seconds shown on the graph, all worms have already passed to the low viscosity media. 30% dextran is used as opposed to pure buffer in order to ease the imaging as the worms swim in this medium but are slower compared to pure buffer. Ten worms were assayed for this experiment.

The point when the droplet fills in the gaps until half the droplet is chosen to be the zero time point and after that 70 seconds lengths of calcium imaging data are analysed. Negative controls are obtained through worms that stay confined within the small droplet as the area surrounding is not filled in order to compensate for photo-bleaching (n=10). Image sequences are imported to ImageJ software^67^ and the fluorescent signal is tracked manually using manual tracking tool^68^. A 30 pixel region of interest around the tracking point is used to extract fluorescence signal where the mean of the signal in this region is measured. This is a bigger area than the cell size, but it is used to capture blurred signals due to quick movement during swimming and also to capture any blurred signals that arise due to loss of lens focus when the worm’s body moved along the z-axis. In some cases, the signal from the region of interest is unreliable due to external interference. For example, a deep omega turn sometimes caused the head dopaminergic neurons, which are also labeled by GCaMP3, to enter the region of interest and interfere with the desired signal. In such cases, the measurement is omitted. Omitted readings are later interpolated using an inpainting function that uses the surrounding time points to create a sparse matrix of spring functions and connections (MATLAB central inpaint nans function). The solution of this function creates a smooth interpolated output (Figure 2). The signal is then smoothed (Moving-average filter, width=10 seconds) to remove movement artefacts. Another region of interest not containing the worm is chosen as a background signal that gets subtracted from the neurons’ signal. The average of 10 seconds before the zero time point is used as a base signal for normalization (Figure 1).

In order to further ensure the accuracy of the measured calcium signal, another negative control from the same worm was measured. This was obtained by measuring the auto-fluorescence signal from the worm’s body near the midpoint. This area is close to the PDE neurons while not having any neurons labelled by GCaMP3 and would serve as a measure of the effect of movement of the worm in the z-axis direction. The mean of the signal from a 50-pixel circular region of interest was measured and normalized using the same method as the neurons’ signals. A bigger region of interest is chosen here because the measured feature is bigger compared to the neurons’ signal and hence the mean of that region is computed, the signal does not need any further normalization.

65% dextran was prepared by dissolving 65 g dextran powder 200,000 (Wako Pure Chemical Industries Ltd., Japan) in 100 mL NGM buffer. Mixing is done using the overhead stirrer DLH (VELP Scientifica, Italy). The mixture is then autoclaved to get rid of air bubbles. 30% dextran was prepared by proportional addition of 65% dextran and NGM buffer and then mixing by sonication. For preparation of NGM buffer, 3 g NaCl + 975 mL H_2_O, 1 mL CaCl_2_, 1 mL MgSO_4_, and 25 mL KH_2_PO_4_ (pH 6.000 adjusted by 5 molar KOH) were mixed after autoclaving to obtain 1 L of the NGM buffer solution. The solution resulting pH was 6.3.

### Imaging

In this experiment, Ca^+2^ imaging was done under Nikon A1R high-speed confocal microscope using its standard widefield fluorescence capability with 10x objective lens (NA=0.25). The samples were illuminated by a mercury lamp (Nikon Intensilight C-HGFIE) with GFP filter. Acquisition is done by Andor Zyla 5.5 high resolution camera (Andor Technology Ltd., US) with Micromanager software^69^. Exposure time was set to 100 ms so the resulting frame rate is 10 FPS and 4 x 4 binning was done on each image so that the final resolution is 640 x 540 pixels. The resulting axial resolution is 8.5 ¼m which is above the typical size of a *c. elegans* neuron cell body thus capturing the signal from the whole cell body with a small tolerance for small z-axis movements. Furthermore, the choice of region of interest size desribed in the previous section, furthr increases the effective slice thickness making measurements even more resistant to z-axis movements. Autotracking was done using Hawkvision movable stage system (Hawkvision Co. Ltd., Japan) equipped with Point Grey GRAS-03K2M-C bright field camera (Point Grey Research Inc, Canada). Tracking software is controlling the stage movements in order to keep the whole body of the worm in the field of view^70^.

## Acknowledgements

I would like to first thank Prof. Ichiro Maruyama for his advice throughout the study. I would also like to thank the Okinawa Institute of Science and Technology, Graduate University for providing the support and funding throughout the whole study period. I am very grateful to Dr. Bernd Kuhn for his advice with fluorescence imaging, and Steve Aird, the technical editor at Okinawa Institute of Science and Technology, for help with editing sections of this manuscript. I would like to thank Prof. Dr. David Biron for sending the dat-1p::GCaMP3. I would like to also thank the current and previous members of the Biology Information Processing unit in the Okinawa Institute of Science and Technology for their help in teaching me different experimental protocols.

## Author contributions statement

The author was responsible for designing the study concept, conducting experiments, data analysis and manuscript writing.

## Additional information

The author declares no conflicting interests. This study was funded through the intramural funding of the Okinawa Institute of Science and Technology.

